# Establishing cell-intrinsic limitations in cell cycle progression controls graft growth and promotes differentiation of pancreatic endocrine cells

**DOI:** 10.1101/2020.03.13.990812

**Authors:** Lina Sui, Yurong Xin, Daniela Georgieva, Giacomo Diedenhofen, Leena Haataja, Qi Su, Yong Wang, Michael Zuccaro, Jinrang Kim, Jiayu Fu, Yuan Xing, Danielle Baum, Robin S. Goland, Jose Oberholzer, Fabrizio Barbetti, Peter Arvan, Sandra Kleiner, Dieter Egli

## Abstract

Limitations in cell proliferation are a key barrier to reprogramming differentiated cells to pluripotent stem cells, and conversely, acquiring these limitations may be important to establish the differentiated state. The pancreas, and beta cells in particular have a low proliferative potential, which limits regeneration, but how these limitations are established is largely unknown. Understanding proliferation potential is important for the safty of cell replacement therapy with cell products made from pluripotent stem cell which have unlimited proliferative potential. Here we test a novel hypothesis, that these limitations are established through limitations in S-phase progression. We used a stem cell-based system to expose differentiating stem cells to small molecules that interfere with cell cycle progression either by inducing G1 arrest, impairing S-phase entry, or S-phase completion. Upon release from these molecules, we determined growth potential, differentiation and function of insulin-producing endocrine cells both *in vitro* and after grafting *in vivo*. We found that the combination of G1 arrest with a compromised ability to complete DNA replication promoted the differentiation of pancreatic progenitor cells towards insulin-producing cells, improved the stability of the differentiated state, and protected mice from diabetes without the formation of cystic growths. Therefore, a compromised ability to enter S-phase and replicate the genome is a functionally important property of pancreatic endocrine differentiation, and can be exploited to generate insulin-producing organoids with predictable growth potential after transplantation.

## Introduction

The unlimited proliferation potential of human pluripotent stem cells is both an opportunity as well as a challenge: it provides an renewable source of cells for cell replacement for degenerative disorders such as diabetes, but it is also a risk for the formation of growths in cell transplantation. Limitations in proliferation potential are established in parallel to limitations in differentiation potential; cells of the adult pancreas have largely stable identities and a very low proliferative and regenerative potential. Limitations in cell proliferation of the pancreas are cell-intrinsic, and established during embryonic expansion of pancreatic progenitors ^1^. Proliferation of beta cells in the developing human pancreas occurs primarily during embryogenesis, and declines after birth, in particular in human beta cells where proliferation in the adult is essentially absent ^2, 3^. During terminal differentiation, many cell types, including neurons and muscle cells exit the cell cycle as they adopt full functionality ^4, 5^. When forced into the cell cycle, adult beta cells and neurons frequently undergo apoptosis ^6, 7^, suggesting a compromised ability to progress and complete S-phase. Whether these limitations in S-phase progression play a functional role in establishing cell-intrinsic limitations in cell proliferation and are important to establish the terminally differentiated state is not known.

Several studies show that cell cycle progression can be disruptive to beta cell function. When beta cells are immortalized to generate proliferating beta cell lines, the differentiated state and function are compromised. For instance, the stable transformed mouse insulinoma cell line MIN6 and the rat insulinoma cell line INS1E progressively develop glucose-independent insulin secretion and begin to express other islet hormones with increasing passage ^8–10^. Overexpression of the pro-proliferation molecules cyclin-dependent kinases in primary rat beta cells increases proliferation, leads to dedifferentiation of primary beta cells and reduces glucose stimulated insulin secretion ^11^. Furthermore, overexpression of the oncogene c-Myc in adult mouse beta cells increases proliferation and beta cells acquire an immature phenotype ^12^. Conversely, removal of immortalizing transgenes in EndoC-βH1 cell line, a proliferative immortalized beta cell line generated from human fetal pancreas, decreases cell proliferation and enhances a number of beta cell specific features such as increased insulin gene expression and content ^13^. Furthermore, several studies show that cell cycle progression is disruptive to the differentiated state in general: cell cycle progression plays a critical role in mediating the transition of a differentiated cell to a pluripotent stem cell ^14–20^. Oncogenic principles disrupting the limitations in cell proliferation of a differentiated cell can facilitate reprogramming, including mutations in p53 ^21, 22^ or Rb ^23^, as well as c-Myc ^24^ or SV40 T-Ag overexpression ^25^. These manipulations directly affect DNA replication and/or impair the exit from the cell cycle in response to genome instability. Hence instability of the genome during DNA replication acts as the primary barrier to somatic cell reprogramming, acting upstream of and prior to barriers affecting changes in transcription ^26^.

We reasoned that the reverse might also apply: that limitations in cell cycle progression can be established through interference with the progression of DNA replication, and that these manipulations promote differentiation to insulin-producing cells from pluripotent stem cells and help to establish cell-intrinsic limitations in growth potential. To test this, we treated pancreatic progenitors with compounds interfering with DNA replication and/or cell cycle progression through different mechanisms and determined beta cell differentiation efficiency, stability of the differentiated state and beta cell function *in vitro* and *in vivo*. Compounds that interfered with G1 to S phase transition, as well as with S phase completion were most effective in increasing differentiation efficiency to insulin-producing cells, resulted in greater stability of the differentiated state. These compounds included the DNA polymerase inhibitor aphidicolin, the antineoplastic agent cisplatin and the topoisomerase inhibitor etoposide. Inducing G1 arrest alone, such as through CDK4 inhibition without compromising transition through S phase was less effective. Upon transplantation, aphidicolin-treated insulin-producing cells demonstrated higher human C-peptide secretion, greater responsiveness to glucose level changes, and protected mice from diabetes without the formation of teratomas or cystic structures. These results demonstrate that limitations in DNA replication link proliferation potential and beta cell identity, which can be exploited to improve graft outcomes in the context of cell replacement for diabetes.

## Results

### Aphidicolin affects DNA replication in a dose-dependent manner in pancreatic progenitors

Aphidicolin (APH) is a DNA polymerase inhibitor which interferes with DNA replication fork progression and has dose-dependent effects on S-phase entry and completion^27 28^. At high concentrations, Aphidicolin inhibits S-phase entry, while at low concentrations, Aphidicolin inhibits S-phase completion and enhances fragility of common fragile sites while S-phase entry is not impaired ^29^. To understand the effect of S-phase entry and completion on pancreatic differentiation from pluripotent stem cells, we exposed cells to different concentration of APH from the pancreatic progenitor cluster (PPC) stage (day 15) to the beta cell (PB) stage (day 27) (Fig. 1A). Cells were exposed to APH from d15-d27 at increasing concentrations from 0.1 μM to 1 μM. We evaluated replication progression by sequentially labeling the cells with thymidine analogs IdU and CIdU for the first hour of APH on day 15 (Fig. 1B). Replication fork speed decreased from 1.6 kb per min in untreated cells to 0.5 kb per minute in cells treated with 0.25 uM or 0.5 uM, and was further reduced at 1 μM APH (Fig. 1C), consistent with earlier studies on replication fork progression at different concentrations of aphidicolin ^28^. The decrease in replication fork progression correlates with cell cycle progression examined on day 18 with 2 hour EdU labeling at day 17 (Fig. 1D, E). Most cells arrested in G1 phase and less than 1% of cells were in S phase when cells were treated with 0.5 μM and 1 μM APH (Fig. 1E). When cells were pulsed with EdU for 2 hours and released to complete the cell cycle (Fig. 1D), cells have an impared replication progression through S phase in the presence of a concentration of 0.25 μM, 0.5 μM and 1 μM APH. Lower concentration of APH allowed S-phase entry and the progression through S phase (Fig. 1F, S1A). Cells in 0.1 μM APH can still progress through S to G1 phase, potentially through compensatory mechanisms.

**Figure 1.**
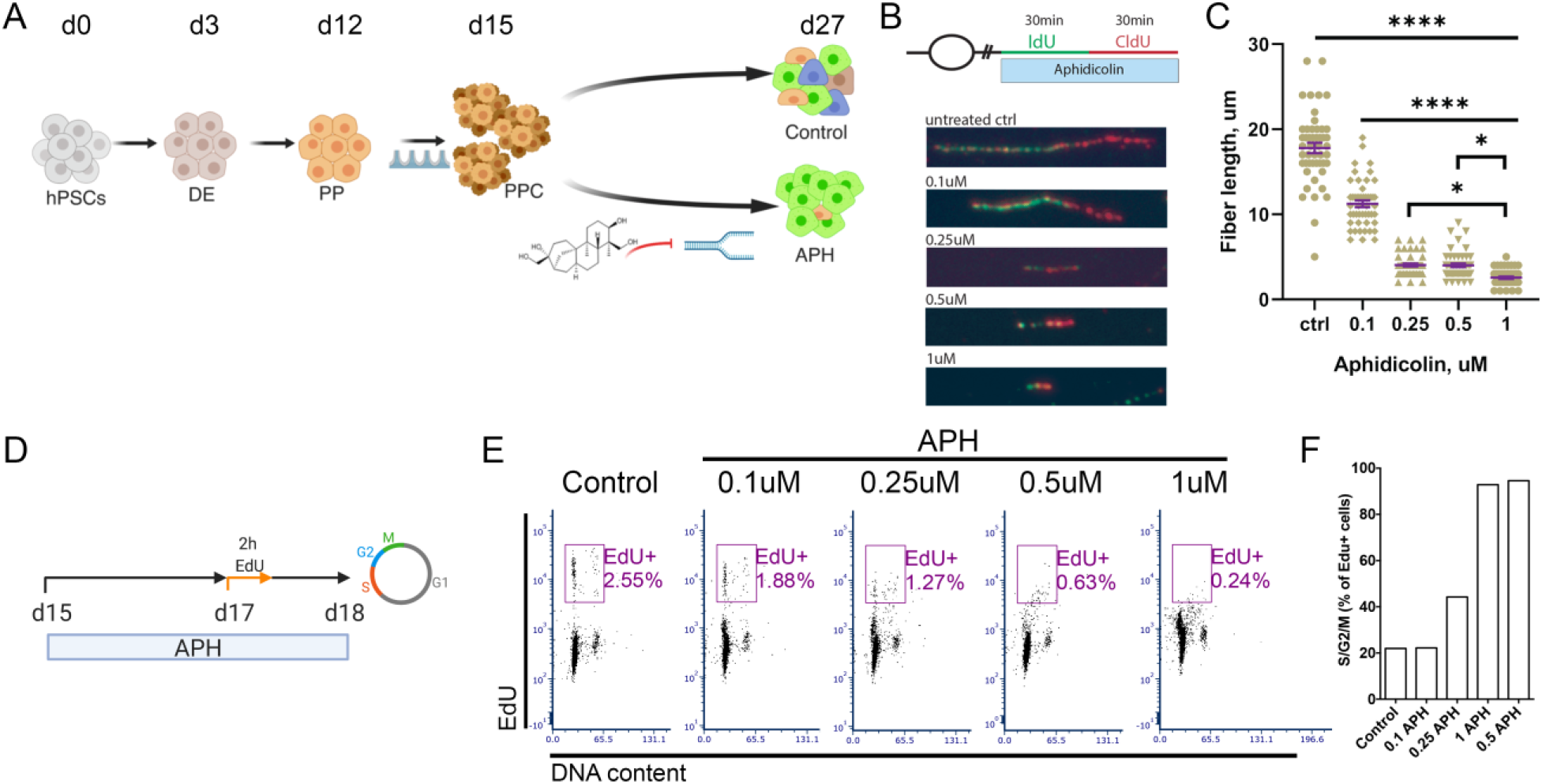
Aphidicolin affects DNA replication progression in a dose-dependent manner in pancreatic progenitors. (**A**) A schematic diagram represents the differentiation of human pluripotent stem cells towards insulin-producing cells with and without APH treatment. hPSCs: human pluripotent stem cells; DE: Definitive endoderm; PP: Pancreatic progenitor; PPC: Pancreatic progenitor cluster. (**B**) DNA replication progression analysis by labeling cells with IdU and CIdU and (**C**) quantification of labeled fiber length. One-way ANOVA with *: p<0.05; ****: p<0.0001. All conditions under **** are significant different from control or 0.1 Μm. (**D**) A schematic diagram indicates the time of APH treatment, EdU incubation and cell cycle analysis. (**E**) Cell cycle profile of cells treated with different concentrations of APH at day 18 after 2 hours of EdU labeling at day 17. (**F**) The percentage of EdU-positive cells in S/G2/M phase and failed to progress to G1 phase were determined at day 18.

### Aphidicolin promotes pancreatic endocrine cell differentiation from stem cells *in vitro*

To determine the effect of aphidicolin on endocrine differentiation, we treated progenitor cells at day 15 till day 27 with indicated concentration of APH and quantified the differentiation efficiency. Aphidicolin concentration of 0.1 μM had no effect on the percentage of pancreatic endocrine cells. Aphidicolin concentrations starting from 0.25 μM resulted in higher differentiation efficiencies than controls, with 1 μM APH giving rise to the highest percentage of C-peptide and NKX6.1-positive cells (Fig. 2A, S1B). 66.3% ± 7.6% C-peptide-positive cells were induced in the condition with 1 μM APH, which was significantly higher than the percentage of C-peptide-posoitve cells generated in control (36.5% ± 14.5%) (Fig. 2B, D). Similar percentage of C-peptide-positive cells coexpressed glucagon (APH: 16.5% ± 4.8%; Control: 15.8% ± 6.5%) or somatostatin (APH: 5.9% ± 2.6%; Control: 8.1% ± 5.5%) in the stem cell-derived endocrine clusters between APH and control (Fig. 2C, D). APH treatment reduced the variability of beta cell differentiation across different experiments with the same embryonic stem cell line. Without APH, the percentage of C-peptide-positive cells ranged widely, from 10%-60% (n=12 independent differentiation experiments). With APH, all (n=12 independent differentiation experiments) cultures contained more than 50% C-peptide-positive cells with the highest over 80% (Fig. 2B). To evaluate if the positive effect of APH was consistent across different genetic backgrounds, we included two iPSC lines with different differentiation potentials, 1018E and 1023A. 1018E was previously identified as a cell line with poor differentiation competence ^30^. The percentage of C-peptide-positive cells was significantly higher after APH treatment in both cell lines (Fig. S1C, D). Remarkably, the poor differentiation potential of 1018E increased to the range of a differentiation competent cell line, from an average of 11% to 38% (Fig. S1C). Therefore, APH increased the purity of stem cell-derived insulin-producing cells after formation of pancreatic progenitors in ESCs and iPSCs of different genetic backgrounds.

**Figure 2.**
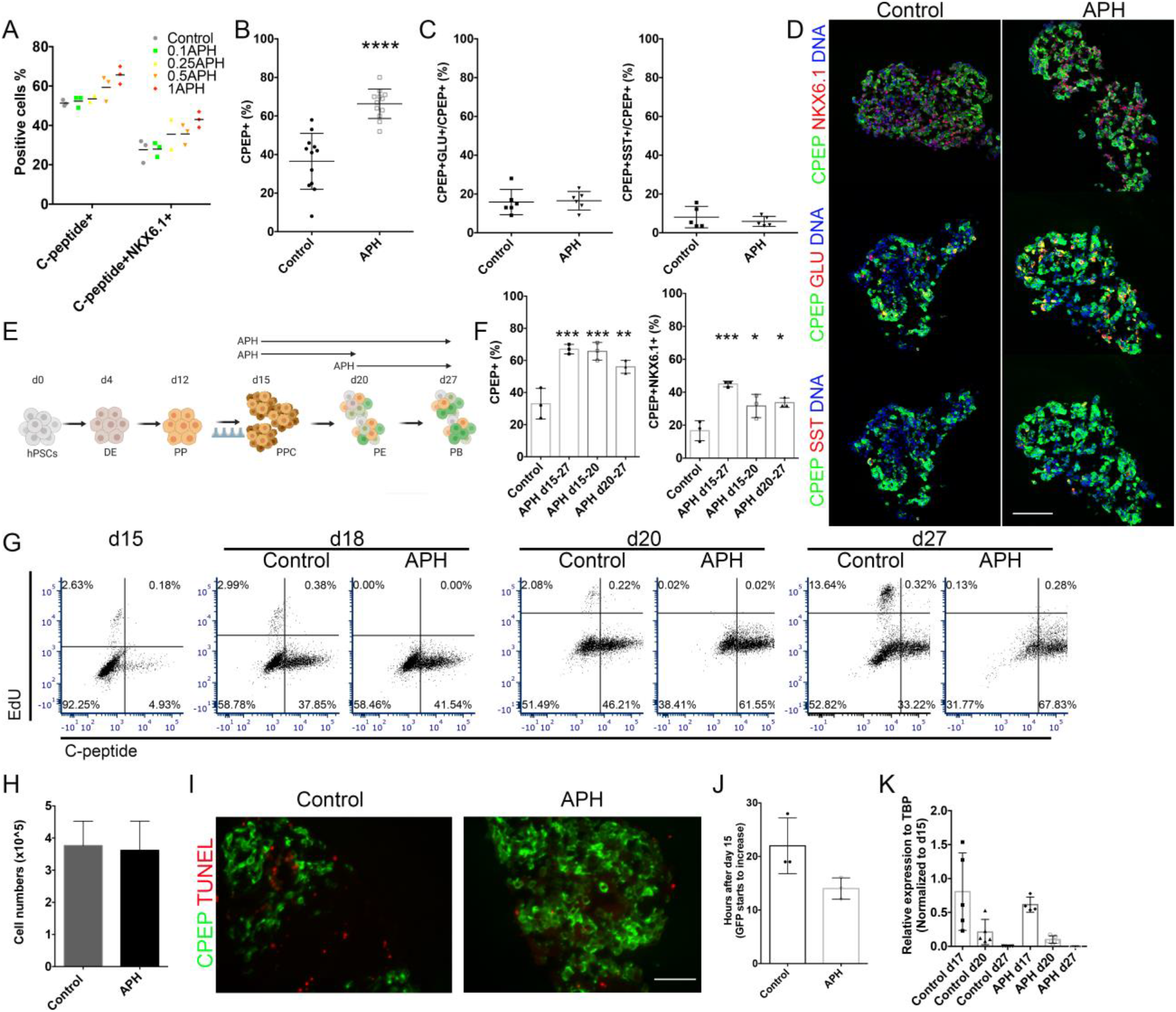
Inhibiting replication fork progression promotes differentiation through cell cycle arrest, not apoptosis. (**A**) Quantification of C-peptide and NKX6.1-positive cells on day 27 treated with different concentrations of APH. (**B**) Quantification of C-peptide-positive cells in control and APH groups. Two-tailed paired t-test with ****: p<0.0001 (**C**) Quantification of C-peptide and glucagon double-positive cells or C-peptide and somatostatin double-positve cells in control and APH groups. (**D**) Immunostaining of frozen section of APH-treated clusters for C-peptide, NKX6.1, glucagon and somatostatin. Scale bar: 100 μm. (**E**) A schematic diagram indicates the duration of APH. hPSCs: human pluripotent stem cells; DE: Definitive endoderm; PP: Pancreatic progenitor; PPC: Pancreatic progenitor cluster; PE: Pancreatic endocrine cells; PB: Pancreatic beta cells. (**F**) Flow cytometry quantification of the percentage of C-peptide-positive and C-peptide and NKX6.1 double-positive cells in the indicated conditions at day 27. One-way ANOVA test with *: p<0.05; **: p<0.01; ***: p<0.001. (**G**) Flow cytometry profile of cell cycle progression at day 15, day 18, day 20 and day 27 of differentiation without and with APH treatment indicated by the percentage of cells stained positive for C-peptide and EdU. (**H**) The total numbers of cells on day 20 of differentiation with and without APH treatment. (**I**) Immunostaining of clusters on d27 with and without APH treatment for TUNEL and C-peptide. Scale bar: 100 μm. (**J**) The time of GFP starts to increase after APH addition at day 15. (**K**) Quantitative RT-PCR determined the NGN3 expression after normalization with TBP and the expression on day 15.

To determine at which stage of differentiation APH acts to promote endocrine differentiation and how it acts, we applied APH at different stages of differentiation from progenitors (PPC) to the beta cell (PB) stage including early stage (d15-d20) during endocrine progenitor development (PPC-PE), late stage (d20-d27) after commitment of endocrine lineages (PE-PB), and throughout the whole duration of the differentiation from pancreatic progenitors to endocrine cells (d15-d27) (PPC-PB), and evaluated the percentage of C-peptide-positive cells as well as C-peptide and NKX6.1 double-positive cells derived at the end of differentiation on day 27 (Fig. 2E). Addition of APH at all indicated stages increased the proportion of C-peptide-positive cells as well as C-peptide and NKX6.1 dual-positive cells (Fig. 2F and Fig. S1E). We profiled cell cycle progression on day 15, day 18, day 20 and day 27 of beta cell differentiation (Fig. 2G). In untreated cells, C-peptide-positive cells started to form on day 15 (~5%) and reached a peak on day 20 in control with ~46% C-peptide-positive cells and ~60% C-peptide-positive cells in APH condition. ~2% of all cells in control and very few of cells, if any in APH condition underwent proliferation during 2 h EdU incubation at each stage before day 20. The total cell number remained the same between two groups (Fig. 2H) and no significant apoptosis was detected after APH treatment either at early stage of day 17 (Fig. S1F) or at late stage of day 27 indicated by TUNEL staining (Fig. 2I). We also traced the expression of insulin-GFP using live-cell imaging starting from day 15 when APH was added. APH-treated progenitor clusters started to express insulin 8 hours earlier than clusters without APH treatment: GFP started to increase at 14.00 ± 1.16 hours in APH and 22.00 ± 3.00 hours in control condition (Fig. 2J and Supplementary Video 1-6), and insulin-GFP glowed brighter in APH-treated clusters compared to control (Supplementary Video 1-6). Therefore, the increased percentage of C-peptide-positive cells from day 15 to day 20 is due to increased differentiation, not primarily by inhibiting the expansion of other cell types and not by inducing cell death of C-peptide-negative cells.

Neurogenin 3 (NGN3) is essential transcription factor expressed in the endocrine progenitors. To test if the increased differentiation efficiency is due to the increased expression of NGN3 in APH treated cells at early stage of differentiation, we checked the transcription levels of *NGN3* in control and in APH treated cells at day 15, d17, d20 and d27. We found that the average expression of *NGN3* was comparable with or without APH treatment, but control cells showed a greater variability in *NGN3* expression levels among each batch of differentiation (n=5) compared to APH treated groups (n=5) (Fig. 2K). This variability might contribute to the variability in the final percentage of C-peptide-positive cells ranging from 10%-60% in controls (Fig. 2B).

From day 20 to day 27, the percentage of proliferating cells increased to ~13% and the percentage of C-peptide-positive cells decreased to 33% in control. The high percentage of C-peptide-positive cells was maintained or increased further from day 20 to day 27 and few proliferating cells were detected in APH condition (Fig. 2G). This suggests that the increased percentage of C-peptide-positive cells in APH is mainly due to the inhibition of proliferation that resumed in control after day 20. Together, adding APH at early and late stage of differentiaton increases the percentage of C-peptide-positive cells through different mechanismes.

### Inhibition of S-phase entry and compromised S-phase completion promote endocrine differentiation

To determine whether the increased differentiation efficiency is specific to polymerase inhibition, or due to its effect on S-phase entry and/or progression, we tested a panel of compounds interfering with either S-phase entry or completion, or both. The CDK4 inhibitor (CDK4i) arrests cells in early G1 phase ^31^. The MCM replicative helicase inhibitor (Ciprofloxacin) and the E2F inhibitor (E2Fi), a transcription factor required for regulating expression of S phase genes ^32^ prevent entry into S phase. Cisplatin (Cis) induces DNA damage by crosslinking DNA, interferes with S-phase progression and arrests cells at G0/G1 phase ^33^; Etoposide (Eto) is a topoisomerase II inhibitor which prevents the unwinding of the DNA helix during replication and transcription and arrests cells mainly in the S and G2 phases ^34^; Aphidicolin (APH) is a DNA polymerase inhibitor which inhibits G1 to S phase transition and S-phase progression in a concentration dependent manner ^27 35^. Other tested compounds include pyridostatin (PDS), a compound promoting the formation of G4 structures to induce replication-fork stalling ^36^, thereby affecting S-phase progression. The percentage of C-peptide-positive cells was increased in all conditions treated with the indicated compounds in comparison to untreated controls (Fig. 3A). A high percentage of insulin-expressing cells indicated by the expression of GFP and evenly distributed in the islet-like clusters were observed with all compounds tested, whereas some parts of the clusters in control remained GFP negative (Fig. 3B). Notably, cells treated with APH and Cis expressed higher levels of GFP compared to cells treated with other compounds and control (Pictures were taken with equal exposure time).

**Figure 3.**
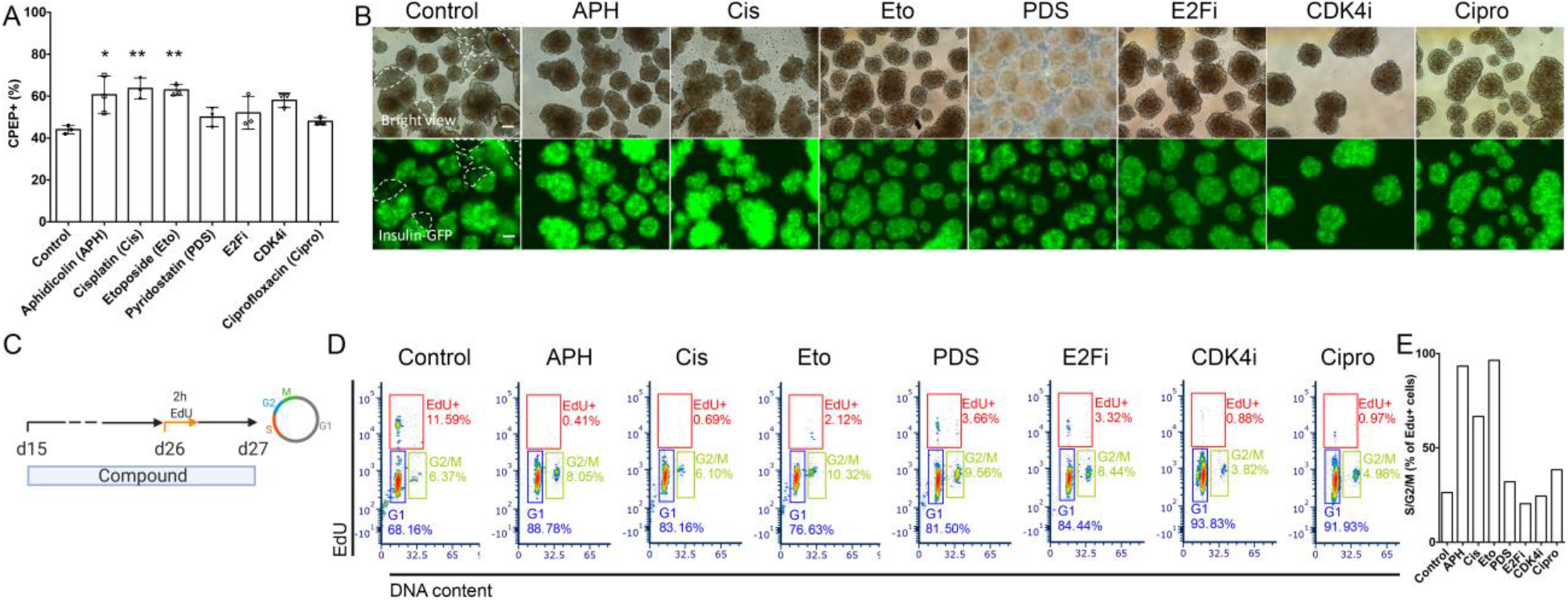
Interference with DNA replication improves differentiation of insulin-producing cells. (**A**) Flow cytometry quantification of C-peptide-positive cells at day 27 with indicated compounds. One-way ANOVA test with *: p<0.05; **: p<0.01. (**B**) Representative bright view and fluorescence picture of stem cell-derived islet clusters after the treatment of indicated compounds. GFP negative parts are circled by dashed white line. Scale bar: 100 μm. Pictures were taken with an OLYMPUS 1X73 fluorescent microscope with equal exposure time (148.1 ms). (**C, D**) EdU pulse and chase experiment to determine S-phase entry and progression. Cells treated with indicated compounds were labeled with EdU for 2 hours on day 26, and analyzed 1 day later for cell cycle distribution. (**E**) Quantification of EdU-positive cells in S/G2/M phase and failed to progress to G1 phase.

We analyzed the cell cycle progression by labeling cells with EdU for 2 hours on day 26 and collected cells for flow cytometry analysis the next day (Fig. 3C). We found that the ability to increase the percentage of insulin-positive cells correlated with an increase in the percentage of G1/G0 cells, a reduction of EdU-positive cells, as well as an increase in the percentage of G2/M cells relative to cells that completed the cell cycle in the presence of the compound and progressed to G1 (Fig. 3D, E and Fig. S2A). Aphidicolin, cisplatin and etoposide result in all three changes to cell cycle progression (Fig. 3D, E and Fig. S2A). Compounds that did not fulfill all three changes, including aphidicolin at low concentration of 0.1μM, were less effective. CDK4 inhibitor and the replication licencing inhibitor ciprofloxacin arrested cells in G1 with comparable efficiency to aphidicolin, but showed only an insignificant increase in endocrine differentiation (Fig. 3A, D). CDK4i had no effect on the progression of replicating cells through G2 to G1 phase (Fig. 3E). E2Fi and PDS reduced the number of cells in S phase, but not to the same extent as other compounds (Fig. 3D). Therefore, induction of G1 arrest and compromised completion of S-phase progression in pancreatic progenitors synergize to improve differentiation to stem cell-derived insulin-producing cells.

### Transcriptome analysis shows an increase in endocrine cells and a decrease in non-endocrine cells

To futher understand the cell composition of stem cell-derived clusters, we performed single-cell RNA sequencing of control and APH-treated cells. We sequenced 16739 cells (8091 cells in control and 8648 cells in APH) on day 27 of differentiation from both conditions (Fig. S3A). We first identified eleven cell populations using cells from both control and APH-treated cells and a large proportion of the cells were endocrine cells (Fig. 4A). We then compared these populations corresponding to primary pancreatic human islets ^37^ (Fig. 4B). Stem cell-derived islet-like clusters contained all endocrine cells identified in primary human islets, including beta, alpha, delta and pancreatic-polypeptide cells. Three stem cell-derived populations (SC-beta 2, SC-alpha, and SC-delta) corresponded most closely to primary human islet beta, alpha, and delta cells (Fig. 4B). 69 genes expressed in pancreatic beta cells and characteristic of mature cells were enriched in SC-beta 2 cluster (including *IAPP*, *SIX2*, *HOPX*, *NEFM*), proinsulin processing and insulin granule exocytosis (*PCSK1*, *CPE*, *PDIA3*, *RAB1A*, *RAB2A*, *RAB3A*, *SCG3*, *VGF*), and metabolism sensing and signaling pathways including (*NUCB2*, *PAM*, *G6PC2*, *PDX1*) (Fig. S3B).

**Figure 4.**
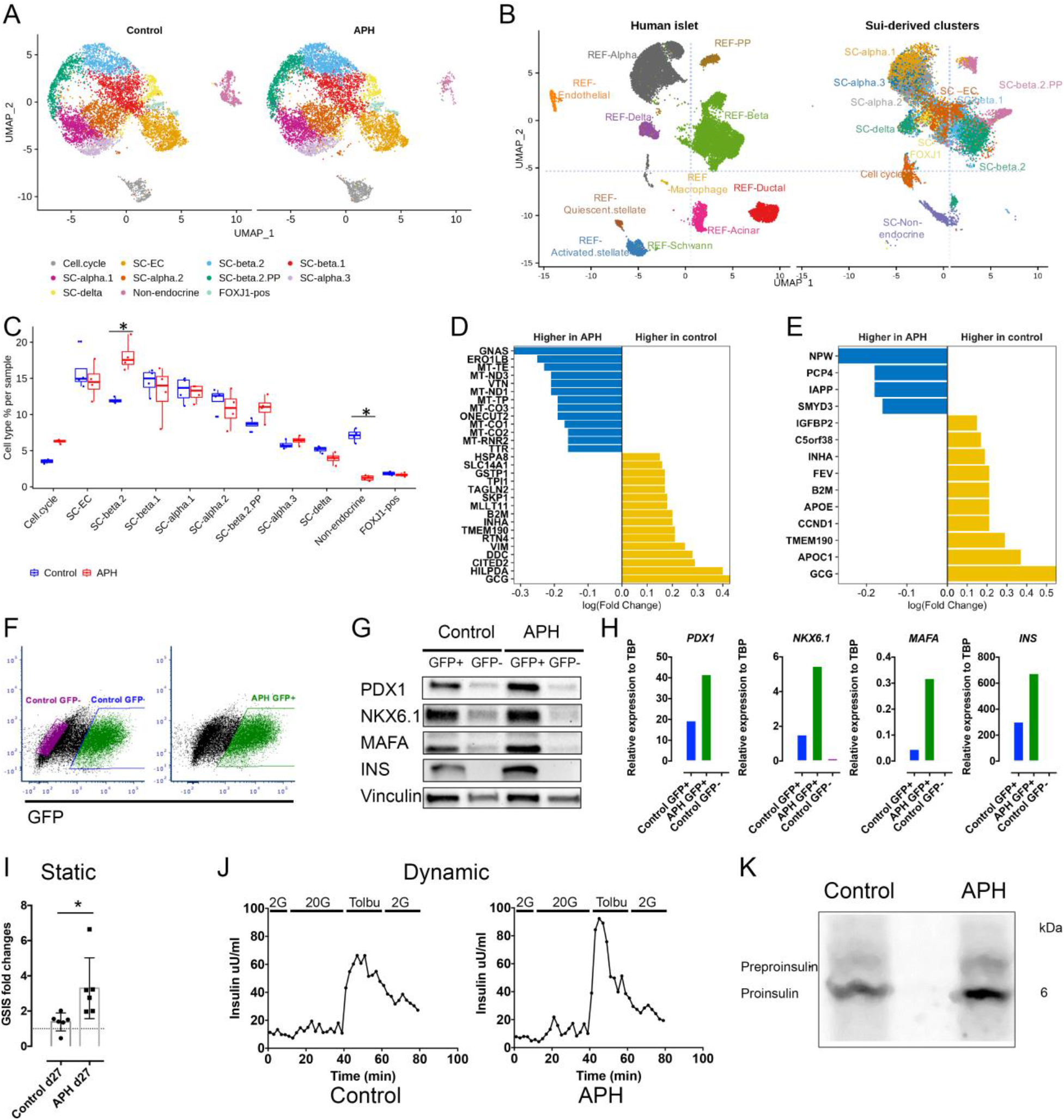
Single cell transcriptome analysis demonstrates the increase of endocrine cells and decrease of non-endocrine cells in APH condition. (**A**) Identified cell populations in stem cell-derived islet cells with and without APH during differentiation. (**B**) Identified cell populations in stem cell-derived islet cells compared to primary human islet cells. (**C**) Quantification of indicated cell populations between control and APH group. *: Wilcox test with p<0.05. (**D**) The upregulated and downregulated genes after cells were treated with APH compared to control in SC-beta 1 cells and (**E**) SC-beta 2 cells. (**F**) Insulin-GFP-positive and negative cells were sorted from control and APH treated group for downstream analysis. (**G**) Protein expressions of sorted cells by Western blot. (**H**) Gene expressions of sorted cells by RT-PCR. (**I**) Static glucose stimulated C-peptide secretion (a stimulation index is determined by the fold changes of C-peptide secretions when control cells and APH-treated cells incubated in the 2 mM and 20 mM glucose). Two-tailed unpaired t-test with *: p<0.05. (**J**) Dynamic analysis of insulin secretion stimulated sequencially by 2mM glucose, 20mM glucose, 150 μM Tolbutamide. (**K**) Proinsulin biosynthesis of an iPSC cell line-derived beta cell clusters with and without APH treatment.

We also identified a group of insulin-expressing cells (SC-beta 1) that failed to overlap with the human primary beta cell cluster (Fig. 4B). Gene Ontology analysis illustrates that genes expressed in SC-beta 1 are enriched in the biological process of glycolytic process and response to hypoxia, whereas genes in SC-beta 2 are highly enriched in the process of hormone transport and secretion (Fig. S3C). Higher expression of lactate dehydrogenase *LDHA* which inhibits mitochondrial activity and lower expression of key beta cell genes was also observed in SC-beta1 (Fig. S3D). This indicates that SC-beta 1 cells are less mature than SC-beta 2 cells. Other endocrine cells included endocrine cells expressing enterochromaffin cell markers (SC-EC) and a cluster of cells with several hormones (and more similar to pancreatic polypeptide cells (SC-beta 2 PP)). Non-endocrine cells partially overlap with acinar cells and duct cells in the human islet. Additional cell types such as endothelial cells or macrophages were only seen in human islets, while stem cell-derived clusters contained additional non-endocrine cells including enterochromaffin cells, FoxJ1 positive cells and cells in the cell cycle (Fig. 4B). In a comparison with single cell RNA sequencing data obtained from a published dataset ^38^, we found a high correlation for each cell cluster (Fig. S4A, B).

We then compared APH-treated and untreated cell populations and quantified their composition. We found a striking reduction in the number of non-endocrine cells in the APH group compared to control and to a published data set ^38^ (Fig. 4C and Fig. S4B). Furthermore, the proportion of insulin expressing cells SC-beta 2 that are most similar to beta cells of primary human islets was significantly increased after APH treatment (Fig. 4C). Changes in the number of cells expressing cell cycle genes (*CDKN1A*, *GADD45A*, *BAX*, *MDM2*, *RPS27L*, *PCNA*, *RRM2*, *CDT1*, and *TYMS*) was also observed, which is likely due to the G1 arrest induced by APH (Detailed data is listed in Fig. 5B). Within SC-beta 1 population, the upregulated genes of APH treated cells included *GNAS*, an important gene for beta cell insulin secretory capacity and function ^39^; *ERO1LB*, a beta cell enriched gene that is involved in insulin processing ^40^; *ONECUT2*, a transcription factor and increased with age in beta cells ^41^; and *TTR*, which has a positive role in glucose stimulated insulin release ^42^. *TPI1*, a gene acting in glycolysis, was decreased in APH-treated cells (Fig. 4D). Within SC-beta 2 cells, upregulated genes included *IAPP* and *PCP4*, a gene involved in Ca^2+^ binding and signaling ^43^. The expression of *FEV*, a signature gene expressed in immature beta cells ^44^, was reduced in APH compared to control (Fig. 4E). We also observed that the cell cycle genes (*CITED2* and *CCND1*) were downregulated compared to control in both SC-beta 1 and SC-beta 2 cells.

**Figure 5.**
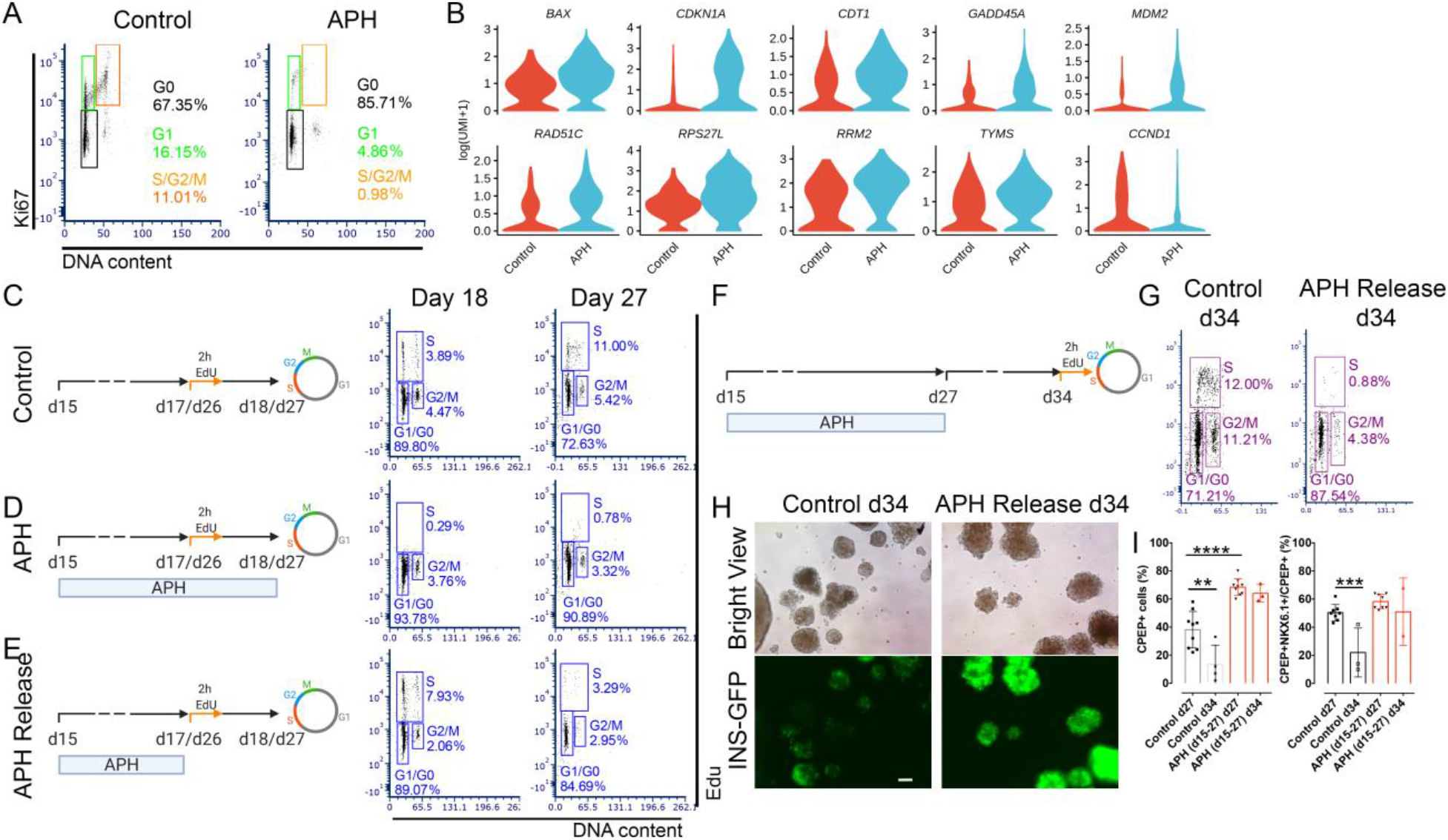
Aphidicolin reduces growth potential and increases beta cell stability by upregulating cell cycle inhibitors. (**A**) Cell cycle profile of untreated and treated cells on day 27 indicated by flow cytometry combined with Hoechst staining for DNA content and Ki67 labeling. (**B**) The differential expression of cell cycle genes between control and APH-treated cells using single-cell RNA sequencing. (**C, D, E**) Schematic diagrams of cell cycle progression experiments with indicated conditions. (**E**) APH-treated cells were released and labeled with EdU for 2 hours either on day 17 or day 26, and analyzed 1 day later (day 18 or day 27) for cell cycle distribution. Cells without APH treatment (**C**) and unreleased from APH (**D**) were analyzed in parallel. (**F**) A schematic diagram represents the time line of APH addition, release from cell culture and cell cycle progression analysis after 2h EdU incubation on day 34. (**G**) Cell cycle distribution of day 34 cells with and without APH from d15 to day 27. (**H**) Microscopic pictures of cells at day 34 after releasing cells from APH from day 27 to day 34. Scale bar: 100 μm. (**I**) Quantification of C-peptide and NKX6.1-positive cells and C-peptide-positive cells at day 27 and day 34 before and after APH releasing. One-way ANOVA test with **: p<0.01; ***: p<0.001; ****: p<0.0001.

To further evaluate beta cell markers in a targeted manner, we isolated insulin-positive cells based on GFP expression (Fig. 4F) and determined the expression of key beta cell genes using Western blot and RT-PCR. We found that protein levels of PDX1, NKX6.1, MAFA and INS insulin were all upregulated in GFP-positive cells isolated from APH treated clusters compared to GFP positive cells in control clusters (Fig. 4G). The higher protein levels correlated with increased transcription levels (Fig. 4H). These data show that aphidicolin-induced cell cycle arrest promotes a gene expression program characteristic of more mature and more functional cells.

To examine the effect of APH on the functionality of insulin-producing cells, static and dynamic glucose stimulated C-peptide secretion was evaluated. In response to elevated glucose, C-peptide secretion was increased 2-6-fold, with an average of 3-fold in a static assay (Fig. 4I). Dynamic perifusion also demonstrated that APH-treated insulin-producing cells showed a relative low secretion of insulin in response to basal glucose and a better response to high glucose and Tolbutamide compared to controls (Fig. 4J). In addition, cells treated with APH produced higher newly-synthesized proinsulin normalized to total protein synthesis (Fig. 4K).

Therefore, cell cycle arrest triggered by aphidicolin not only reduces the proportion of non-endocrine cells, but also increases differentiation efficiency, and results in more mature insulin-producing cells with improved functional properties.

### APH limits growth potential and stabilizes beta cell identity by upregulating cell cycle inhibitors in vitro

To determine whether aphidicolin treatment has lasting effects on cell cycle progression, we examined the expression of cell cycle markers at the end of treatment on day 27. 85% of cells treated with APH versus 67% of cells in control conditions were found to exit the cell cycle to G0 (Fig. 5A). 27% of control cells were in the cell cycle, as indicated by the expression of Ki67 and DNA content. About 5% of APH-treated cells expressed Ki67 in G1, and very few of the cells were in S phase (Fig. 5A). We also examined the cell cycle gene expression in insulin-expressing cells by sorting out the GFP-positive cells from cell clusters. We found that the expression of *CDKN1A* (a cell cycle progression inhibitor) was upregulated, and the expression of *CCND1* (cyclin D1) and *CDK4* (both involved in G1-phase progression) were downregulated in APH-treated insulin-positive cells (Fig. 5B and Fig. S5A). These results demonstrate that APH promotes G1/G0 arrest and induce S and G2/M arrest in pancreatic endocrine progenitors.

A subset of cells appeared competent of DNA replication after aphidicolin treatment. In Ki67-positive cells, single cell RNA sequencing showed upregulation of genes involved in the P53 signaling pathway (*CDKN1A*, *GADD45A*, *BAX*, *MDM2*, *RPS27L*, *PCNA*, *RRM2*) were significantly upregulated in APH-treated cells. These genes mediate G1 cell cycle arrest, or respond to difficulties in DNA replication progression. *CDKN1A* and *GADD45A* are able to arrest cells either in G1/S or G2/M ^45^. *CDT1*, a replication licensing gene (stable in G1 and degraded in S phase), as well as the replication clamp protein *PCNA* were also upregulated, indicating that cells were arrested in late G1 (Fig. 5B). *CDT1* and *PCNA* upregulation appears to be a compensatory response to enable S-phase progression and attempt to rescue S-phase in the presence of APH.

To explore the consequences of transient APH treatment on proliferation potential, we exposed pancreatic progenitors to APH and released them at different time points (Fig. 5C, D, E). On day 17 of differentiation, 2 days after APH treatment, APH was removed from culture and cells were incubated with EdU for 2 hours and collected the next day (day 18). Over 90% of cells were arrested in G1/G0 and very few cells went through S phase when APH was present (Fig. 5D). Upon removal of APH, 8% of cells resumed proliferation, as indicated by EdU staining, more than in control (4%) (Fig. 5E). When cells were treated for 12 days and released from APH on day 26, 3% of cells were EdU positive on day 1 after releasing (day 27), while in controls 11% were proliferating (Fig. 5C, E). Therefore, while short term exposure induced synchronization at the G1/S phase transition, long term exposure resulted in G0 arrest, while ~3% were in G1 (Fig. 5A) and capable of re-entering S phase (Fig. 5E).

To test the stability of G0/G1 arrest, we continued culturing cells after release from APH for 7 days till day 34 and labelled with EdU on day 34 for 2 hours (Fig. 5F). The percentage of EdU-positive cells significantly reduced to 0.88%, whereas control cells continued proliferating at the same rate as on day 27 (Fig. 5F, G). Thus, growth potential was greatly reduced when cells were exposed to APH for a long term period (12 days), and the vast majority of cells had entered a stable G0 state.

To explore if the lasting effect of APH on cell cycle progression contributes to maintain beta cell identity, insulin-producing cells were cultured for additional 7 days till day 34 upon removal of inhibitors on day 27. In untreated control cells and CDK4i-treated cells, the insulin-GFP expression was lost while it remained high in APH-, Cis-, and Eto-treated cells on day 34 (7 days after release) (Fig. 5H and Fig. S5B). In addition, the percentage of C-peptide and NKX6.1 double-positive cells and the total of C-peptide-positive cells was still high on day 34 in cells treated with APH, Cis, and Eto, respectively, whereas the percentage of C-peptide-positive cells was significantly reduced in the control and the CDK4i group. (Fig. 5I, Fig. S5C). Cells treated with low dose of APH (0.1 uM) failed to maintain the stability of C-peptide-positive cells (Fig. S5D). These data show that transient treatment with high dose of APH (>=0.25 uM) (and other inhibitors of DNA replication) resulted in more stable beta cell identity and that the stability of insulin-expressing endocrine cells subsequently becomes independent of the compounds.

### Aphidicolin treated cells show reduces growth potential *in vivo* in a dose-dependent manner

To determine long term effects of APH on growth potential, we removed APH on day 27 and monitored the graft growth after transplantation *in vivo*. We grafted mice with an equal number (~2 millions) of APH-treated cells or with untreated cells. APH was removed permanently on day 27 before transplantation. Graft growth was evaluated by monitoring a luciferase reporter using *in vivo* imaging. After 2 weeks of transplantation, the graft size of APH mice was small, while controls grafted with the same number of cells were modestly larger (Fig. 6A). 11 weeks later, 4/7 control mice displayed large growths, whereas none of the mice transplanted with APH treated cells did (9/9 numbers) (Fig. 6A). The different growth trend of grafted cells between control and APH-treated cells was evident in the bioluminescence intensity (Fig. 6B). At 22 weeks of engraftment, the size of graft in the APH group was on average 2.6-fold larger than that at 2 weeks, while size of the control group increased on average 53-fold (Fig. 6C). Graft growth occurred in controls even from cultures with very high differentiation efficiency (>60%). Even in mice with the smallest growths of control cells, the cystic structures still formed in 3/3 mice (Fig. 6B indicated by blue lines). No cysts were observed in mice grafted with APH-treated cells in 9/9 mice (Fig. 6D).

**Figure 6.**
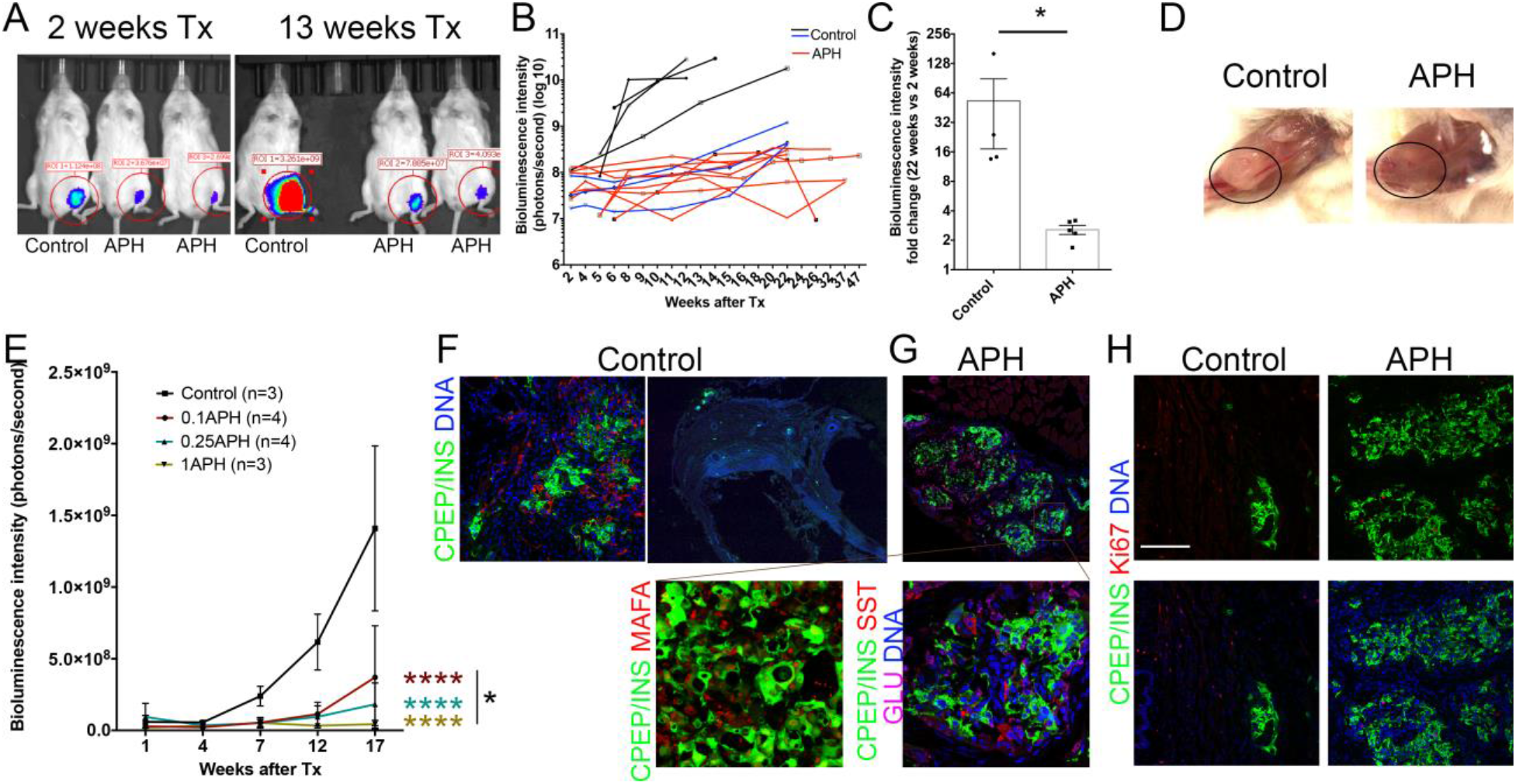
APH treatment prevents teratoma formation and reduces growth potential *in vivo*. (**A**) *In vivo* imaging of mice transplanted with control cells and APH treated cells at 2 weeks and 13 weeks post transplantation. (**B**) Growth of grafted cells in mice after transplantation indicated by the bioluminescence intensity. (**C**) Fold changes of grafted cell growth in control mice and APH mice at 22 weeks of engraftment compared to 2 weeks of transplantation. *: Mann-Whitney test with p<0.05. (**D**) Pictures of grafted cells with equal high differentiation efficiency in control group and APH group at 36 weeks of transplantation. (**E**) Growth of grafted cells in mice after transplantation of cells pretreated with different concentration of APH indicated by the bioluminescence intensity. Two-way ANOVA with *: p<0.05 (0.1APH vs 1APH); ****: p<0.0001 (0.1, 0.25, 1APH vs control). (**F**, **G**) Cell composition determined by immunostaining in mice transplanted with control cells (**F**) and APH-treated cells (**G**). Pictures in bottom panel are close-ups for APH picture. (**H**) Cell proliferation determined by staining grafted cells in control (left) and APH (right) for Ki67. Scale bar: 100 μm.

To determine if APH regulates cell growth in a dose-dependent manner, we transplanted cells pretreated with 0 μM, 0.1 μM, 0.25 μM and 1 μM of APH into mice and monitored the graft growth with bioluminescence intensity for 17 weeks. To ensure a comparable number of cells engrafted, we started the monitoring within a week of transplantation. Similar bioluminescence intensity of grafted cells was observed among 4 groups at 1 week of transplantation and difference in graft size was apparent at 7 weeks of transplantation. Significant differences in grafte size were detected between each group at 12 weeks and 17 weeks after transplantation. As expected, cells without APH treatment underwent substantial proliferation with time and cells treated with APH proliferated less compared to control and in a dose-dependent manner (Fig. 6E).

Grafts were isolated from mice for examination. APH graft were composed of islet-like structures and showed monohormonal cells positive for insulin, for glucagon or for somatostatin (Fig. 6G). Insulin-expressing cells stained positive for MAFA, a master regulator for beta cell maturation (Fig. 6G). In mice grafted with untreated control cells, the graft still contained groups of C-peptide-positive cells but developed several large cystic structures (Fig. 6F) with a large number of cells positive for Ki67 (Fig. 6H). Grafts derived from APH-treated cells had lower, but higher than zero Ki67-positive cells (Fig. 6H), demonstrating that aphidicolin had altered growth potential, and not merely abolished proliferative capacity. We also transplanted 1159 iPSC-derived clusters into NSG mice to determine if growth control by APH also applied to iPS cells. No cysts were observed in mice grafted with APH-treated iPSC-derived clusters (3/3), whereas cysts were formed in mice transplanted with non-treated cells (2/2) (Fig. S6A). Therefore, aphidicolin controlled cell growth in a dose-dependent manner and prevented teratomas and cystic growth after transplantation.

### Cells treated with APH secrete C-peptide and efficiently protect mice from diabetes

To test the ability of APH-treated islet-like clusters to regulate blood glucose levels, we monitored C-peptide and blood glucose levels. After transplantation in immunodeficient mice, the mice transplanted with APH-treated cells trended to have higher human C-peptide starting from 2 weeks after engraftment, compared to controls (Fig. 7A). The increase was statistically significant at 6 weeks after transplantation (Fig. 7A and Fig. S6B, C). Secretion of human C-peptide in mice was downregulated when mice were fasted and increased after glucose injection (Fig. 7B), indicating the engrafted insulin-producing cells were able to respond to changes in blood glucose levels.

**Figure 7.**
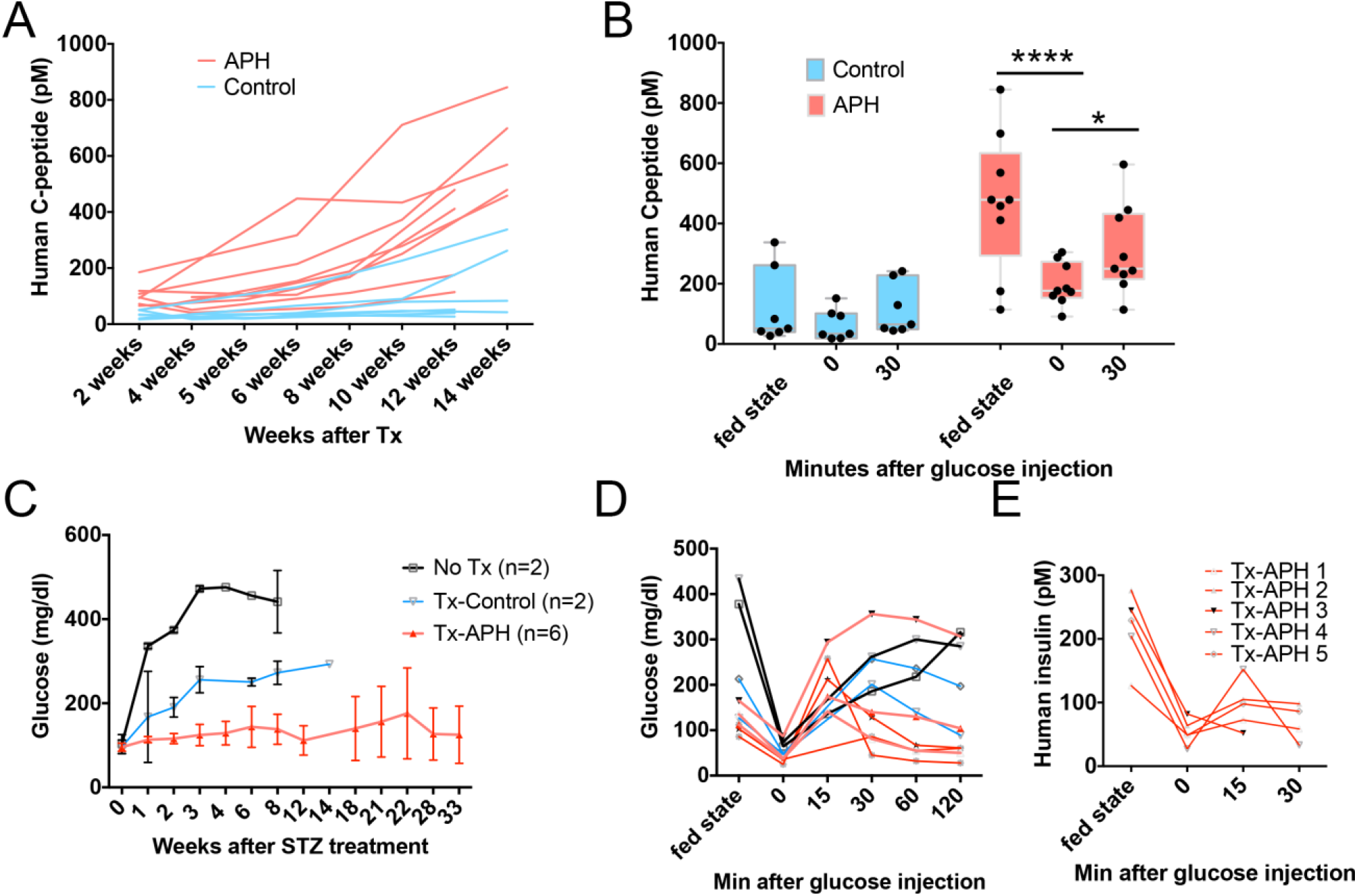
APH treatment prevents diabetes of a mouse model. (**A**) Human C-peptide serum concentration in mice at different time points after transplantation with cells treated with APH (APH) and control cells (MEL1) at fed state. (**B**) Human C-peptide serum concentration in mice at 12-14 weeks after transplantation with cells treated with APH (APH) and control cells (MEL1) at fed state, fasting and 30 min after glucose injection. Two-way ANOVA with *: p<0.05; ****: p<0.0001. (**C**) Blood glucose levels of STZ-treated mice without transplantation (no Tx) (n=2), transplanted with control cells (Tx-Control) (n=2) and with APH treated cells (Tx-APH) (n=6). (**D**) Glucose tolerance test of STZ-treated mice in fed state, fasting state and 15-120 min after glucose injection. (**E**) Serum human insulin concentrations of STZ-treated mice transplanted with APH-treated cells at fed state, at fasting and 30 min after glucose injection.

The ability of APH-treated cells to protect mice from diabetes was determined after eliminating endogenous mouse beta cells with streptozotocin (STZ). STZ ablates mouse beta cells but is not toxic to human beta cells at the concentrations used ^30^. 15 weeks after transplantation, mice were treated with STZ, blood glucose levels were monitored and grafted insulin-producing cells were challenged with high glucose to check their function. Blood glucose levels remained in the normal range (Fig. 7C). Mice were tolerant to glucose and normalized blood glucose levels within 15 min of glucose injection (Fig. 7D). Secretion of human C-peptide and insulin decreased after fasting and increased after glucose injection (Fig. 7E and Fig. S6D). Therefore, APH-treated cells controlled graft growth while protecting mice from diabetes more efficient than non-treated cells.

## Discussion

Substantial progress has been made to differentiate pluripotent stem cells to insulin-producing cells ^46–49^. In common to these studies is the activation or inhibition of developmental signaling pathways to promote differentiation to specific cell types. Here, we test the duplication of the genome as a developmentally relevant target to induce the differentiation and maturation of stem cell-derived insulin-producing cells and to establish cell type specific limitaitons in cell proliferation. Unlike the modulation of signaling pathways that can have complex effects on both gene expression and cell cycle progression, the use of aphidicolin is highly specific in targeting the duplication of the genome by inhibiting DNA polymerase in a dose-dependent manner. Aphidicolin affects both the progression from G1 to S phase, while also affecting DNA replication completion, in particular at common fragile sites ^29^.

A number of compounds tested here arrested pancreatic progenitors in G1, but only DNA replication inhibitors aphidicolin, cisplatin and etoposide significantly increased the production of insulin-producing cells. The noticible difference of DNA replication inhibitors from other cell cycle inhibitors is the compromised cell cycle progression from S to G1 phase. S phase cells in the presence of DNA replication inhibitors failed to complete the cell cycle and arrested during the progression to G1 (Fig. 1E and S1A). The combination of G1 arrest with reduced S phase progression and completion of the cell cycle was most effective in promoting differentiation and stability of the differentiated state. Inhibition of DNA replication reduced variation in differentiation efficiency both in different experiments with the same cell line, as well as with different cell lines. Stem cell lines that were previously demonstrated as differentiation incompetent ^30^ also showed markedly improved differentiation efficiency. Single cell RNA-sequencing data revealed that the number of mature beta cells (SC-beta 2) that are transcriptionally more similar to human primary beta cells was increased. Aphidicolin-treated cells showed upregulated expression of genes in metabolic signaling, insulin processing and release and downregulated expression of genes in cell cycle progression and glycolysis. Furthermore, aphidicolin treatment greatly decreased the number of non-endocrine cells, both in comparison to our controls, as well as compared to another published study ^38^ (Fig. S5AB). Though the resulting cells are similar to pancreatic islets, differences to adult pancreatic cells remain.

Our studies demonstrate an antagonistic relationship between cell proliferation and the commitment of pancreatic progenitors to the endocrine lineage during differentiation from pluripotent stem cells. The study of DNA replication adds a new perspective to the existing literature on cell cycle progression and DNA replication in the pancreatic lineage. Reduced S phase competence is established naturally in pancreatic differentiation, though with slower developmental kinetics than induced by aphidicolin or antineoplastic agents: human beta cells are not only arrested in G1, but they are also compromised in their ability to complete S phase. Forced cell cycle entry of beta cells can induce apoptosis during S phase ^7^. In the brain, replication incompetence is complete: neuronal cells that are forced to re-enter a cell cycle through inhibition of Rb will die and degenerate rather than divide and grow ^6, 50^. Therefore, mechanisms that impair S phase entry and S-phase completion may be an important and fundamental principle of terminal differentiation in several organs, in particular in the brain, in muscle cells and in the pancreas. S-phase completion and S-phase re-entry are functionally linked, as entry into quiescence in G1 is determined by successful completion of the previous S-phase ^51, 52^. Other studies have focused on developmental signals and the activity of transcription factors, such as NGN3 in understanding growth and terminal differentiation of the pancreas^53^. However, how NGN3 and other transcription factors affect cell cycle progression and DNA replication remains to be further investigated. Transcription factors may modulate S-phase entry and progression not only through the expression of gene products, but also by altering origin activity and/or affecting transcription-replication conflicts.

Our findings are relevant to defining growth potential of cell replacement products for the treatment of diabetes. In this and previous studies, a proliferative non-endocrine population in untreated controls can contribute to growths after transplantation ^30, 54^. Several strategies have been developed to reduce the risk of such outcomes. A flow-cytometry based method can be used to purify stem cell-derived insulin-producing cells labeled with a fluorescent reporter to prevent teratomas ^48^. Furthermore, genetically engineering a suicide gene in a stem cell line can efficiently kill non-insulin-producing cells upon drug administration at the end of differentiation ^55^. The rationale for the methods developed here is based on the biology of the mature beta cell: cell cycle exit and a compromised ability to undergo S phase. Through the use of small molecules promoting cell cycle exit, growth after transplantation is controlled and teratomas and cysts are avoided. This method does not require transgenes or flow cytometry and is readily transferable between different cell lines. Remarkably, aphidicolin-treated cells showed consistent graft size up to one year after transplantation. Therefore, we show that growth limitations can be modulated by specific and transient interference with DNA replication. Though we do not know how closely aphidicolin mimics developmental processes, it is of interest to note that neurons exciting the cell cycle incur frequent breaks and copy number changes ^56, 57^, often in late replicating regions that are limiting to replication completion and targeted by aphidicolin treatment ^29^.

In conclusion, we demonstrate an active role of limitations in DNA replication in the stable commitment to an endocrine cell fate during differentiation of pluripotent stem cells. Understanding the molecular mechanisms that limit replication potential in the pancreas is central to regenerative medicine, morphogenesis during development, as well as for the understanding of pancreatic cancer.

## Material and Methods

### Human pluripotent stem cell culture

Human pluripotent stem cells were cultured and maintained on feeder-free plates with StemFlex Medium (Cat. No. A3349401, Thermo Fisher Scientific) as described ^58^. 4 cell lines were involved in this study as shown in Supplementary Table 1: MEL1 is human embryonic stem cell line ^59^; 1023A is a human induced pluripotent stem cell line derived by reprogramming a skin fibroblast biopsied from a healthy control; 1018E is a human induced pluripotent stem cell line reprogrammed from a skin fibroblast biopsied from a female type 1 diabetes patient ^30^. 1159 is a human induced pluripotent stem cell line reprogrammed from a skin fibroblast biopsied from a female healthy control ^30^. All human subjects research was reviewed and approved by the Columbia University Institutional Review Board, and the Columbia University Embryonic Stem Cell Committee.

### Insulin-producing cell differentiation from human pluripotent stem cells

Insulin-producing cells differentiated from human pluripotent stem cell lines using published protocol (Sui et al., 2018b). Briefly, cells were cultured for 4 days using STEMdiff™ Definitive Endoderm Differentiation Kit (Cat. No. 05110, STEMCELL Technologies) for definitive endoderm induction. Media was changed to RPMI 1640 plus GlutaMAX (Cat. No. 61870-127, Life Technology) + 1% (v/v) Penicillin-Streptomycin (PS) (Cat. No. 15070-063, Thermo Fisher Scientific) + 1% (v/v) B-27 Serum-Free Supplement (50x) (Cat. No. 17504044, Life Technology) + 50 ng/ml FGF7 (Cat. No. 251-KG, R&D System) for 2 days. From day 6 to day 8, media was changed to DMEM plus GlutaMax (DMEM) (Cat. No. 10569-044, Life Technology) with 1% (v/v) PS + 1% (v/v) B-27 + 0.25 μM KAAD-Cyclopamine (Cat. No. 04-0028, Stemgent) + 2 μM Retinoic acid (Cat. No. 04-0021, Stemgent) + 0.25 μM LDN193189 (Cat. No. 04-0074, Stemgent). To generate pancreatic progenitors, cells were refreshed in DMEM + 1% (v/v) PS + 1% (v/v) B-27 + 50 ng/ml EGF (Cat. No. 236-EG, R&D System) + 25 ng/ml FGF7 for 4 days. At day12, cells were dissociated into single cells and clustered in AggreWell 400 6-well plates (Cat. No. 34425, STEMCELL Technologies) with DMEM + 1% (v/v) PS + 1% (v/v) B-27 + 1 μM ALK5 inhibitor (Stemgent, cat. no. 04-0015) + 10 μg/ml Heparin (Sigma-Aldrich, cat. no. H3149) + 25 ng/ml FGF7 + 10 μM Y-27632, ROCK inhibitor. At day 13, newly formed clusters were transferred into Low-attachment 6-well plates (Cat. No. 07-200-601, Thermo Fisher Scientific) and cultured with RPMI 1640 plus GlutaMAX + 1% (v/v) PS + 1% (v/v) B-27 + 1 μM thyroid hormone (T3) (Cat. No. T6397, Sigma) + 10 μM ALK5 inhibitor + 10 μM zinc sulfate (Sigma-Aldrich, cat. no. Z4750) + 10 μg/ml heparin +100 nM gamma-secretase inhibitor (DBZ) (EMD Millipore, cat. no. 565789) + 10 μM Y-27632, ROCK inhibitor for 7 days. From day 20 to day 27, clusters were cultured in RPMI 1640 plus GlutaMAX + 1% (v/v) PS + 1% (v/v) B-27 + 10% (v/v) fetal bovine serum (Atlanta Biologicals, cat. no. S11150) + 10 μM zinc sulfate + 10 μg/ml heparin.

### Treatment with DNA replication inhibitors

The DNA polymerase inhibitor aphidicolin (Cat. No. A0781, Sigma-Aldrich) was added with indicated concentrations at specified time point from day 15 to day 27 or as shown in Figures. The concentrations used in this study are 0.1 μM, 0.25 μM, 0.5 μM and 1 μM. 2.5 μM Cisplatin (Cat. No. 232120, EMD Millipore), 2 μM Etoposide (Cat. No. E55500, Research Products International), 10 μM Pyridostatin (Cat. No. S7444, Selleck Chemicals), 10 μM E2F inhibitor (Cat. No. 324461 EMD Millipore), 2 μM Palbociclib Isethionate (CDK4i) (Cat. No. S1579, Selleck Chemicals), and 100 μM Ciprofloxacin (Cat. No. S2027 Selleck Chemicals) were added from day 15 to day 27. The concentrations of compounds were determined based on the survival of cells.

### Replication progression analysis

Cell clusters at day 15 of differentiation were incubated sequentially with 25 uM IdU and 25 uM CIdU for 30 min each in the presence of aphidicolin at 0.1 uM, 0.25 uM, 0.5 uM or 1 uM concertation. Cell clusters were then collected and dissociated into single cells and DNA fibers were stretched and stained for IdU and CIdU as described in Terret et al., 2009. Briefly, drops of cell suspension in PBS were spotted on glass slides and lysed with 0.5% SDS, 200mM Tris pH 7.4, 50mM EDTA. Fibers were then stretched by tilting the slides at a 10-15° angle and fixed with methanol: acetic acid = 3:1. Staining was performed with the following antibodies- anti- BrdU/CldU (Biorad # OBT0030) and anti-BrdU/IdU (BD # 347580). Stained DNA fibers were imaged with the 100x objective of an Olympus inverted fluorescence microscope IX73 and measured with Olympus cellSens analysis software. 1μm was considered 2.6kb as described ^60^.

### Cell cycle progression analysis

Cell clusters at day 17 or day 26 in the indicated conditions were incubated with EdU for 2 hours and then washed twice to remove EdU. Cell clusters were continuously cultured in the medium with or without indicated compounds and collected at the next day and dissociated for flow cytometric analysis. EdU was stained by Click-iT™ EdU Alexa Fluor™ 555 Imaging Kit (Cat. No. C10338, Fisher Scientific) and followed by Hoechst 33342 staining. Flow cytometry was performed to determine the number of cells in each phase of cell cycle.

### Flow cytometry

The beta cell clusters were dissociated using TrypLE™ Express into single cells. Cells were fixed with 4% PFA for 10 min and permeabilizated with cold methanol for 10 min at −20 °C. Primary antibodies were added at a dilution of 1:100 in autoMACs Rinsing Solution (Cat. No. 130-091-222, Miltenyi Biotec) containing 0.5% BSA at 4 °C for 1 hour. Secondary antibodies were added at a dilution of 1:500 at room temperature for 1 hour. Hoechst 33342 (Cat. H3570, Thermo Fisher Scientific) was added 1:2000 dilution together with 2^nd^ antibodies if DNA content was determined. The cells were filtered with BD Falcon 12 x 75 mm tube with a cell strainer cap (Cat. No. 352235, BD) and subsequently analyzed by a flow cytometer. Data was analyzed with the FCS Express 6 Flow software. Negative controls were included by only adding secondary antibodies.

### Gene expression analysis using RT-PCR

Insulin-GFP-positive and -negative cells were sorted with flow cytometry. Total RNA was extracted from the sorted cells from different conditions using a RNeasy Mini Kit (Cat. No. 74106, QIAGEN). The total RNA was reverse transcribed into cDNA using iScript™ Reverse Transcription Supermix (Cat. No. 170-8841, Bio-Rad), and the cDNA was then used as template with SsoFast™ EvaGreen® Supermix (Cat. No. 172-5202, Bio-Rad) for quantitative real-time PCR. The primers used for PCR are listed in Supplementary Table 2.

### Immunocytochemistry

Clusters at d27 were collected and fixed with 4% paraformaldehyde (PFA) at room temperature for 10 min. Grafts were taken from the mice and fixed with 4% PFA at room temperature for 1 hour. The following steps were performed according to the published method (Sui et al., 2018b). Primary antibodies are listed in Supplementary Table 3 and secondary antibodies are listed in Supplementary Table 4. Pictures were taken with an OLYMPUS 1X73 fluorescent microscope or ZEISS LSM 710 confocal microscope.

### *In vitro* fluorescence live cell imaging

Live-cell imaging experiments were performed with EVOS FL Auto (Life Technologies). Clusters were plated onto Geltrex in 96-well plate format and cultured for 3 days in EVOS FL Auto chamber. 3 individual clusters were monitored for indicated conditions. Temperature was set to 37 °C, CO_2_ was set to 5%. Time-lapse images were taken with 470/22 light cube (Life Technologies) every 30 minutes and compiled to create videos.

### Single-cell RNA sequencing and read mapping

Single cells were suspended in PBS+0.04% BSA. Totalseq-A anti-human hashtag antibodies (BioLegend) were used for cell hashing. Each sample was individually stained with one of the hashtag antibodies and washed three times. Eight samples were pooled at equal concentrations, and the pool was loaded into the 10X Chromium instrument at 32000 cells per lane. Single-cell RNA-seq libraries were prepared using Chromium Single Cell 3′ Reagent Kits v2 (10X Genomics). Hashtag libraries were generated as described in ^61^. Sequencing was performed on Illumina NextSeq500. Sequence alignment and expression quantification were performed using Cell Ranger Single-Cell Software Suite (10X Genomics, v2). Reads were aligned to the B37.3 Human Genome assembly and UCSC gene model.

### Single-cell data analysis

Cell-hashing tags were demultiplexed using HTODemux function (Seurat v3). Empty droplets and doublets were excluded. Cells were excluded by following criteria: 1) total UMI > or < 3 fold of median absolute deviation (MAD); 2) detected genes > or < 3 fold of MAD; 3) detected genes in the first of the bimodal distribution (classified by mclust); 4) mitochondrial gene ratio > 0.15. Cells from two experiment groups (Control and APH) were integrated and clustered to identify cell types and subpopulations (Seurat v3). The differentially expressed genes between the two groups were defined with 1.28-fold change and 0.05 adjusted p-value.

### Gene ontology analysis

The functional enrichment analysis was performed using g:Profiler (version e97_eg44_p13_d22abce) with g:SCS multiple testing correction method applying significance threshold of 0.01 ^62^.

### Static glucose stimulated C-peptide secretion

Krebs Ringer buffer (KRB) was prepared by addition of 129 mM NaCl, 4.8 mM KCl, 2.5 mM CaCl_2_, 1.2 mM MgSO_4_, 1 mM Na_2_HPO_4_, 1.2 mM KH_2_PO_4_, 5 mM NaHCO_3_, 10 mM HEPES and 0.1% BSA in deionized water and was sterilized with a 0.22 μm filter. 2 mM and 20 mM glucose solution were prepared in KRB for low glucose and high glucose challenge of sc-beta cell clusters. 10-20 sc-beta cell clusters (~5×10^5^ cells) were collected from control and APH treated conditions and pre-incubated in 500 μl low glucose solution for 1 hour. Clusters were then washed once with low glucose solution and subsequently incubated in 200 μl of low glucose and then high glucose solution for 30 min. 130 μl supernatant from each condition was collected. Supernatants containing C-peptide, secreted in low and high glucose conditions were processed using Mercodia Ultrasensitive Human C-peptide ELISA kit. Fold changes of C-peptide secretion before and after glucose stimulation were calculated.

### Dynamic glucose stimulated insulin secretion

Microfluidic-based perifusion system was used to determine glucose stimulated insulin secretion. This experiment was conducted according to previously described methods (Adewola et al., 2010; Mohammed et al., 2009). 100 clusters from each group were perfused sequentially with KRB containing 2 mM glucose (10 min), 20 mM glucose (30 min), 150 μM Tolbutamide (20 min) and 2 mM glucose (20 min). Perifusate samples were collected at the outlet at flow rate of 200 μl/min and every 2 min samples were used to determine insulin concentration by Mercodia Ultrasensitive Human Insulin ELISA kit.

### Proinsulin biosynthesis

The iPSC cells were recovered overnight in RPMI 1640 plus GlutaMAX + 1% (v/v) PS + 1% (v/v) B-27 + 10% (v/v) fetal bovine serum + 10 μM zinc sulfate + 10 μg/ml heparin with and without APH. The iPSC clusters were picked in 16mM glucose containing RPMI-1640 medium lacking Cys/Met (Cat. No. R7513, Sigma-Aldrich), washed twice with the same medium and pulse labeled for 15 min with ^35^S-Cys/Met (300 μCi, Cat. No. NEG072007MC, Perkin Elmer). After pulse, cells were washed once with cold PBS containing 20 mmol/L N-ethylmaleimide (NEM, Cat. No. E3876, Sigma-Aldrich) and then lysed in radioimmunoprecipitation assay buffer (25 mmol/L Tris (Cat. No. L-27436, Fisher), 1% Triton X-100 (Cat. No. 93443, Sigma), 0.2% deoxycholic acid (Cat. No. BP349, Fisher), 0.1% SDS (Cat. No. 46-040-CI, Corning), 10 mmol/L EDTA (Cat. No. V4231 Promega), and 100 mmol/L NaCl (Cat. No. S9888, Sigma) plus 2mmol/L NEM and a protease inhibitor cocktail (Cat. No. 11836153001, Roche). Cell lysates, normalized to tricholoroacetic acid (Cat. No. BP555-1, Fisher) precipitable counts, were precleared with pansorbin (Cat. No. 507861, Millipore) and immunoprecipitated with anti-proinsulin (Abmart) and protein G-agarose (Cat. No. X1196, Exalpha) overnight at 4ΰC. Immunoprecipitates were analyzed by nonreducing Tris-tricine-urea-SDS PAGE, with phosphorimaging.

### Transplantation and *in vivo* assay

8-10 weeks old male immuno-compromised mice (NOD.Cg-Prkdcscid Il2rgtm1Wjl/SzJ (NSG) from Jackson laboratories, Cat. No. 005557) were used for transplantation. For intra leg muscle transplantation, 1-2 million cells were collected and transferred to a tube with 50 μl Matrigel. They were injected in the leg muscle with 21Gauge needle. The human C-peptide levels in mouse serum were measured every two weeks in the fed state. Intraperitoneal glucose tolerance test was performed by fasting overnight and injecting 2 g/kg D-glucose solution at 2 weeks after mouse beta cells were ablated with one high dose (150 mg/kg) of streptozotocin (Cat. No. S0130-1G, Sigma-Aldrich). Blood was collected in heparin-coated tubes at fed state, at fasting state and 30 min after glucose injection. Plasma was collected by centrifuging tubes at 2000 g for 15 min at 4 °C. The supernatants were collected for C-peptide and insulin detection with Mercodia M-Plex™ ARRAY Chemiluminescent Mercodia Beta Kit. Blood glucose levels were measured at fed state, at fasting state and every 30 min after glucose injection for 2 hours. All animal protocols were approved by the Institutional Animal Care and Use Committee of Columbia University.

### *In vivo* imaging

Mice were injected with 150 mg/kg D-luciferin (Cat. LUCK-2G, Gold Biotechnology). Subsequently, mice were anesthetized and placed in IVIS Spectrum Optical Imaging System (PerkinElmer) 10 min after injection. Pictures were taken analyzed with Living Image Software. Graft size was determined by bioluminescence intensity.

### Western blotting

The sorted GFP-positive and -negative cells with and without APH treatment were lysed with RIPA buffer. Protein was quantified with Pierce BCA protein assay kit. 20 μg of protein was loaded on 4-20% Mini-PROTEAN TGS Precast Protein Gels (Cat. No. 4561094, Bio-Rad) from each condition and ran at 100 V for 1 hour followed by transferring the protein on a PVDF membrane. The membrane was blocked with 3% BSA in TBST for 1h at RT and incubated with primary antibodies diluted in TBST with 3% BSA overnight at −4 °C on a shaker. After washing with TBST, the membrane was incubated with secondary antibodies diluted in TBST with 3% non-fat dry milk powder for 1 h at room temperature. The image was taken with the ChemiDoc Imaging Systems (Bio-Rad).

### Statistical analysis

Statistical tests performed for specific data sets are described in the corresponding figure legends. Data was analyzed by an unpaired or paired t test and by one-way or two-way ANOVA followed by Tukey’s multiple comparison test (GraphPad Prism 6, GraphPad Software, Inc., La Jolla, CA) and plotted as mean ± standard deviation (SD). The differences observed were considered statistically significant at the 5% level and were displayed on figures as follows: *p<0.05, **p<0.01, ***p<0.001, ****p<0.0001.

### Data availability

Single-cell transcriptome data is deposited in Gene Expression Omnibus GEO: GSE139949. All other data supporting the findings of this study are available from the corresponding author on reasonable request.

### Code availability

All code information used for single cell RNA sequencing analysis will be available from the corresponding author on reasonable request.

## Author Contribution

L.S. conceived and designed the study, performed experiments, analyzed data, and wrote the manuscript with input from all authors; Y.X. performed single cell RNA sequencing analysis; J.D. performed western blot and RT-PCR; D.G. performed DNA fiber analysis; L.H. and P.A. performed proinsulin biosynthesis analysis; Y.W. and Y.X. performed microfluidic perifusion and data analysis; M.Z. performed live imaging of stem cells-derived insulin-positve cells; J.F. assisted with GSIS, *in vivo* imaging and ipGTT; D.B. assisted with drug testing; Q.S. optimized single cell RNA sequencing analysis; J.K. and S.K. dissociated beta cell clusters and prepared cells for single cell RNA sequencing; R.G. coordinated human subject research; S.K. discussed RNA seq result. D.E. conceived the study, contributed to experimental design, data interpretation and manuscript writing. D.E., R.G., P.A., S.K., F.B and J.O. provided supervision.

## Acknowledgments

We acknowledge Hang Song, Christina Adler and Evangelia Malahias from Regeneron Pharmaceuticals, Inc. for generating and preparing hash tag single cell libraries and NGS sequencing. This research was supported by Russell Berrie Foundation program in cellular therapies of diabetes, Columbia DERC grant (to L.S), NYSTEM IDEA award C029552, Leona and Harry Helmsley Charitable Trust (2015PG-T1D069), Regeneron Pharmaceuticals, HIRN UC4 DK104207 and RO1 DK103585, American Diabetes Association ADA 1-16-ICTS-029 (to D.E) and the Weezie Foundation. NIH DK48280 and NIH DK111174 (to L.H and P.A), the JDRF Microfluidic-Based Functional Facility at University of Virginia (J.O), and the Chicago Diabetes Project (CDP). D.B. is supported by the National Center for Advancing Translational Sciences, National Institutes of Health, through Grant Number TL1TR001875. F.B. is supported by a Scientific Career Development Grant by European Society for Paediatric Endocrinology (ESPE). These studies used the resources of the Diabetes and Endocrinology Research Center Flow Core Facility funded in part through Center Grant 5P30DK063608. INS^GFP/W^-hES cells is a gift from Dr. Edouard G. Stanley. We thank Dr. Rudolph L. Leibel and Dr. Megan Sykes for helpful discussions. Research reported in this publication was performed in the CCTI Flow Cytometry Core, supported in part by the Office of the Director, National Institutes of Health under awards S10RR027050 and S10OD020056. Lina Sui is a Russell Berrie Foundation Scholar.

